# Functional characterization of QT interval associated *SCN5A* enhancer variants identify combined additive effects

**DOI:** 10.1101/2024.03.11.584440

**Authors:** Lavanya Gunamalai, Parul Singh, Brian Berg, Leilei Shi, Ernesto Sanchez, Alexa Smith, Ghislain Breton, Mark T Bedford, Darius Balciunas, Ashish Kapoor

**Affiliations:** Institute of Molecular Medicine, University of Texas Health Science Center at Houston, Houston, Texas 77030; Department of Integrative Biology and Pharmacology, McGovern Medical School, University of Texas Health Science Center at Houston, Houston, Texas 77030; Department of Epigenetics and Molecular Carcinogenesis, University of Texas MD Anderson Cancer Center, Houston, Texas 77030; Department of Biology, College of Science and Technology, Temple University, Philadelphia, Pennsylvania 19122

**Keywords:** *SCN5A*, enhancer, variant, QT interval, additive, transcription factor

## Abstract

Several empirical and theoretical studies suggest presence of multiple enhancers per gene that collectively regulate gene expression, and that common sequence variation impacting on the activities of these enhancers is a major source of inter-individual variability in gene expression. However, for vast majority of genes, enhancers and the underlying regulatory variation remains unknown. Even for the genes with well-characterized enhancers, the nature of the combined effects from multiple enhancers and their variants, when known, on gene expression regulation remains unexplored. Here, we have evaluated the combined effects from five *SCN5A* enhancers and their regulatory variants that are known to collectively correlate with *SCN5A* cardiac expression and underlie QT interval association in the general population. Using small deletions centered at the regulatory variants in episomal reporter assays in a mouse cardiomyocyte cell line we demonstrate that the variants and their flanking sequences play critical role in individual enhancer activities, likely being a transcription factor (TF) binding site. By performing oligonucleotide-based pulldown assays on predicted TFs we identify the TFs likely driving allele-specific enhancer activities. Using all 32 possible allelic synthetic constructs in reporter assays, representing the five biallelic enhancers in tandem in their genomic order, we demonstrate combined additive effects on overall enhancer activities. Using transient enhancer assays in developing zebrafish embryos we demonstrate the four out the five enhancer elements act as enhancers *in vivo*. Together, these studies extend the previous findings to uncover the TFs driving the enhancer activities of QT interval associated *SCN5A* regulatory variants, reveal the additive effects from allelic combinations of these regulatory variants, and prove their potential to act as enhancers *in vivo*.

## Introduction

Among various gene expression regulation mechanisms, *cis*-regulation is the most common (1), and large-scale epigenomic studies have uncovered multiple putative *cis*-regulatory enhancers per gene (2). Along with driving precise spatiotemporal control of gene expression (3), multiple enhancers per gene likely act as buffer against disruptive sequence variation within enhancers (4). Furthermore, presence of noncoding variants-based multiple independent association signals within genome-wide association studies (GWAS) loci (5) and the use of multiple putative regulatory variants in gene expression prediction models for most genes in humans (6) and model organisms (3, 7), also suggest that multiple enhancer variants collectively regulate gene expression as we shown by us (8, 9) and others (10). Although enhancer cooperation has been studied to some extent at the level of individual genes (11, 12), the nature of these combined effects from multiple enhancers and their variants remains functionally unexplored for most genes, especially for genes implicated for common diseases and traits by GWAS. Here, we characterize five *SCN5A* enhancer variants, identified earlier to be underlying the electrocardiographic QT interval GWAS locus on chromosome 3p22.2 (8, 13), to assess their combined effects on reporter gene expression.

Electrocardiographic QT interval, a clinically relevant heritable quantitative trait, measures the time taken by cardiac ventricles to de- and re-polarize in every heartbeat (14). QT interval varies across individuals, is correlated with age, sex and heart rate, and has a heritability of 35% (15). Prolongation or shortening of QT interval, owing to underlying pathology, genetic causes or adverse drug reactions, is associated with increased risk for cardiac arrythmias and sudden cardiac death, both in the general population (16) and in the clinical extremes (17). With rare, high-penetrance deleterious coding variants in *SCN5A* leading to Mendelian long QT syndrome-3 (LQT3) (17) and common noncoding variants at the *SCN5A* locus associated with the trait in the general population (13, 18), *SCN5A*, encoding the voltage-gated sodium channel alpha subunit, has been established as a major genetic regulator of QT interval variation. Following the mapping of QT interval GWAS locus at *SCN5A*, we had performed an unbiased *in vitro* screen for enhancers overlapping all common variants associated with the trait and uncovered five high-confidence causal enhancer variants, one each for the five independent association signals within the GWAS locus, which collectively correlated with *SCN5A* cardiac expression and thereby explained the QT interval associations (8).

Here, we report functional characterization of these five *SCN5A* enhancers and their variants. We use constructs bearing variant-centered small deletions to demonstrate that the enhancer activity of these elements in reporter assays is dependent on the variants and their flanking sequences. We use oligonucleotide (olig0)-based pulldowns of transcription factors (TF) predicted to bind variants and their flanking sequences to identify the putative TFs driving allele-specific enhancer activities. We use synthetic constructs representing all 32 allelic combinations of the five biallelic enhancers in tandem in reporter assays and identify additive effects. And, we use transient zebrafish enhancer assays to demonstrate that the majority of these elements act as enhancers *in vivo*. Together, this study uncovers the putative TFs driving allele-specific activities of the *SCN5A* enhancer variants, their combined additive effects on reporter expression and confirms their potential to act as enhancers *in vivo*.

## Methods

### Cloning of variant-centered deletion allele for in vitro reporter assays

±5-base deletion alleles (11-base deletion) centered on variants of interest were generated using a four primers-based two-step PCR approach with the internal reverse and internal forward primers containing the desired 11-base deletion in their overlapping region (see Dataset S1 for primer sequences). Deletion alleles, like the previously cloned reference and alternate alleles (8), were cloned upstream of a minimal promoter driven firefly luciferase reporter gene in the pGL4.23 vector (Promega). Existing plasmids constructs of variant-centered test elements for the five variants of interest (see Dataset S1 for genomic regions) cloned in the pGL4.23 vector (Promega) were used as the initial template in PCR. PCR was performed in a 25 µL volume containing 10 ng of plasmid DNA or 5 ng each of upstream and downstream amplicons, 0.2 mM (each) dNTP, 0.5 µM forward primer, 0.5 µM reverse primer, 0.5 unit of Q5 High-Fidelity DNA Polymerase (New England Biolabs), and 1× Q5 buffer (New England Biolabs). Thermal cycling was performed as follows: initial denaturation 98 °C for 30 s, 15 cycles of denaturation 98 °C for 10 s, annealing 65 °C for 20s, extension 72 °C for 15 s, and final extension of 72 °C for 2 min. An aliquot (10%) of PCR product was visualized by 2% agarose TAE (Tris base, acetic acid and EDTA) gel electrophoresis. PCR amplicons were purified using DNA Clean & Concentrator-5 kit (Zymo Research) following the manufacturer’s recommendations. Purified PCR products (deletion-bearing, second-step amplicon) and the pGL4.23 vector (Promega) were double digested with NheI-HF (New England Biolabs) and XhoI (New England Biolabs), gel-purified using Zymoclean Gel DNA Recovery kit (Zymo Research), and ligated using T4 DNA Ligase (New England Biolabs), all following manufacturers’ recommendations. Chemically-competent *E. coli* (DH5α) cells (5o µL) were transformed with 2 µL of the ligation reaction and 1/10^th^ of the outgrowth was spread on selective medium [LB agar + ampicillin (5o µg/mL)], and incubated overnight at 37 °C. Bacterial colonies were inoculated in LB broth + ampicillin, incubated at 37 °C and 225 rpm overnight, and then harvested for glycerol stock and plasmid preparations (Zymo Research). Positive clones were identified by restriction digestion (NheI-HF and XhoI) and Sanger sequencing of plasmid DNA using pGL4.23 vector backbone primers (Dataset S1).

### Cloning of allelic combinations across the five candidate elements for in vitro reporter assays

Synthetic constructs representing the 32 allelic combinations at the five enhancer variants of interest (five variants each with two alleles; 2^5^ combinations) were cloned in pGL4.23 using HiFi DNA Assembly technology (New England Biolabs). Variant-centered, reference and alternate-allele bearing elements were first amplified to generate overlapping fragments in their genomic order, with the first and the last elements overlapping with KpnI and XhoI digested ends of pGL4.23 vector, respectively, and then assembled and cloned using HiFi DNA Assembly master mix (see Dataset S1 for primers). PCR amplification, gel check, PCR purification, vector preparation and identification of positive clones were performed as described above.

### Luciferase reporter assays in HL1 cells

Mouse cardiomyocyte HL1 cells (a gift from William C. Claycomb (deceased), Louisiana State University) were maintained in Claycomb medium (Sigma) as described earlier (19). Reporter constructs were transfected into HL1 cells, grown in 24-well plates at ∼80% confluency, using FuGENE HD Transfection Reagent (Promega) following the manufacturer’s recommendations. pRLSV40 (Promega), expressing *Renilla* luciferase, was co-transfected to normalize for transfection efficiency. Forty-eight hours after transfection, cells were harvested and lysed, and firefly and *Renilla* luciferase activities were measured on a microplate reader (Tecan) using the Dual-Luciferase Reporter Assay System (Promega) following the manufacturer’s recommendations. Reporter assays were performed for each construct in four replicates. For each measurement, the observed firefly luciferase reading was divided by the observed *Renilla* luciferase reading to get relative firefly activity. Relative firefly activity was divided by the average relative firefly activity from empty vector (pGL4.23), and then averaged across replicates, to obtain the mean normalized reporter activity for each test construct. The significance of allelic difference was evaluated by comparing mean normalized reported activities at allelic pairs using the Student’s *t* test. All *P* values reported are two-sided.

### Cloning of variant-centered test amplicons for transient enhancer assays in zebrafish

Plasmid pGG4, a zebrafish transient enhancer assay vector capable of both Tol2 transposase mediated random integration and PhiC31 integrase mediated site-specific integration in a compatible host (20), was first modified using a polylinker DNA (HindIII-XhoI-EcoRV-EcoRI-SphI-BamHI-XbaI; Dataset S1) to include a multiple cloning site (MCS) between HindIII and XbaI restriction sites and generate pGG4-MCS vector. For each of the five variants of interest, reference, alternate and deletion-alleles centered elements were amplified (see Dataset S1 for primers) from the pGL4.23 backbone plasmids and cloned as HindIII-XbaI fragments in pGG4-MCS. PCR amplification and cloning were performed as described above. Positive clones were identified by restriction digestion (HindIII-HF and XbaI) and Sanger sequencing of plasmid DNA using pGG4-MCS vector backbone primers (Dataset S1). Repeated attempts of cloning rs6801957 centered elements in pGG4-MCS failed, therefore, these elements were cloned as SalI-BamHI fragment in pT2-mcfos-eGFP vector (21), another zebrafish transient enhancer assay vector for Tol2 transposase mediated random integration.

### Transient enhancer assays in zebrafish

All protocols for zebrafish care and use were reviewed and approved by the Institutional Animal Care and Use Committees at Temple University (TU) and University of Texas Health Science Center at Houston (UTHealth), and zebrafish were cared for by standard methods (22). Tol2 transposase mRNA was *in vitro* transcribed using the mMESSAGE mMACHINE T3 Transcription kit (ThermoFisher Scientific). For injections, 300 ng of reporter plasmid, 300 ng of transposase mRNA, and phenol red (0.1% final concentration) were combined in a 20 uL volume and injected into wildtype AB strain embryos (∼3 nL/embryo) at 1-2 cell stage by a pico-injector (Harvard Instruments). At 24h post fertilization (hpf), embryos were moved to phenylthiourea-containing egg water to reduce pigmentation that helps in better imaging of fluorescent reporter signal. Embryos were imaged 72hpf.

### In silico prediction of TF binding

Based on the human reference genome, for each of the five enhancer variants, two 21-base long sequences centered at the variant, corresponding to the reference and alternate alleles were identified as input sequence (Dataset S2). We scanned these 10 sequences using FIMO (23) against the JASPAR 2022 CORE vertebrates (non-redundant) database (24) with default settings and match output filtered for *P*<1×10^-4^. The database contains 841 non-redundant motifs.

### Cloning of DNA-binding domains of candidate TFs as GST-fusion proteins

DNA-binding domains of candidate TFs were identified using PhosphoSitePlus (25). Gene synthesis service at Biomatik (Ontario, Canada) was used to clone DNA-binding domains of a subset of the predicted TFs (Dataset S2) as bacterial codon-optimized open reading frames in pGEX-4T1 (Amersham) to generate N-terminal Glutathione S-transferase (GST) fusion proteins. Positive clones were identified by restriction digestion (BamHI-HF and XhoI) and Sanger sequencing of plasmid DNA using pGEX-4T1 vector backbone primers (Dataset S1).

### Expression and purification of GST-fusion proteins

GST-fusion constructs in pGEX-4T1 backbone were transformed in chemically-competent BL21 strain of *E. coli* for protein expression. Protein induction by isopropyl-1-thio-β-D-galactopyranoside (IPTG), bacterial lysis by sonication, affinity purification using glutathione sepharose matrix (Amersham), and elution of GST-fusion protein using reduced glutathione buffer were performed following the manufacturer’s recommendations and as described earlier (26).

### DNA-TF interactions

DNA oligo-based pulldown assays were performed to assess interactions between enhancer variant and its flanking sequence and predicted TFs as described earlier (27). Oligos with the following stem-loop structure were designed and HPLC purified (Integrated DNA Technologies): 5’ end Biotin-tri ethylene glycol (TEG) modification, bases encompassing the variant site and its flanking sequence predicted to bind specific TFs, (T)_4_ that forms the loop, and the reverse complement (Dataset S1). Oligos (50µM) were self-annealed by denaturation and slow cooling in oligo annealing buffer (10mM Tris pH 8.0, 10mM NaCl, 1mM EDTA) to form the double-stranded stem-loops. Streptavidin agarose beads (ThermoFisher Scientific) were washed three times in cold binding buffer (50mM Tris pH 7.5, 250-350mM NaCl, 0.05% IGEPAL CA-630) to remove storage buffer. For oligo pulldown assay, 2µg of GST-fusion protein was incubated with 4µg biotinylated annealed oligo in 200µL cold binding buffer overnight at 4°C with rotation, followed by addition of 30µL of 50% Streptavidin agarose beads slurry, and incubated for another hour at 4°C with rotation. Following centrifugation at 500×g for 3 min and removal of supernatant, beads were washed three times with 1ml cold binding buffer (4°C with rotation for 5 min followed by 500×g spin for 3 min for each wash). Captured proteins were denatured using equal volume of 2× SDS-PAGE sample loading buffer and analyzed by 10% SDS-PAGE and Coomassie blue staining following standard methods (28). For comparisons, 2µg of GST-fusion protein alone was used as an input and a pulldown without biotinylated oligo (GST-fusion protein + Streptavidin agarose beads) was used as a negative control.

## Results

### Deletion of QT interval associated variants and their flanking sequence impacts the activities of the five SCN5A enhancers

In an earlier screen for *cis*-regulatory enhancers centered on QT interval associated common variants at the *SCN5A* GWAS locus using *in vitro* luciferase reporter assays in mouse cardiomyocyte HL1 cells (19), we had identified five enhancers with significant difference in allelic activities between the reference and alternate alleles at rs7373779, rs41312411, rs11710077, rs13097780 and rs6801957 (8). Given that the activities of these five enhancers were variant allele dependent, we evaluated if the variant along with immediate flanking sequence were critical for enhancer activity. At each of the five enhancers, deletion alleles, bearing ±5 base-deletion centered on the variant (11-base deletion), were generated by PCR (Dataset S1) and, like the reference and alternate allele constructs, were cloned upstream of a minimal promoter-driven firefly luciferase reporter gene in pGL4.23 vector. Luciferase reporter assays were performed in mouse cardiomyocyte HL1 cells using reference, alternate and deletion allele constructs for each of the five enhancer elements. The enhancer activities observed in deletion alleles at four out of the five enhancer elements were significantly reduced compared to reference and/or alternate alleles (Figure 1). At rs7373779, rs41312411 and rs13097780 centered enhancers, deletion alleles had reduced activities compared to both reference and alternate alleles, suggesting that the 11-base deletions removed binding sites for TFs critical for enhancer function. At rs11710077 centered enhancer element, where the reference and alternate alleles had the largest fold-change (over 2-fold) in enhancer activities among the five enhancers, the deletion allele partially rescued the loss of function observed at the alternate allele-bearing element. This suggests that rs11710077 alternate allele, but not the reference allele, along with the flanking sequence is likely bound by a TF that represses the enhancer. At rs6801957 centered enhancer, the deletion allele completely rescued the loss of function observed at the reference allele-bearing element, suggesting that rs6801957 reference allele, but not the alternate allele, along with the flanking sequence is likely bound by a TF that represses the enhancer. In summary, enhancer activities from deletion alleles at all five enhancer elements were significantly different from reference and/or alternate alleles suggesting that the variant sites and their flanking sequences are critical for enhancer function.

**Figure 1:**
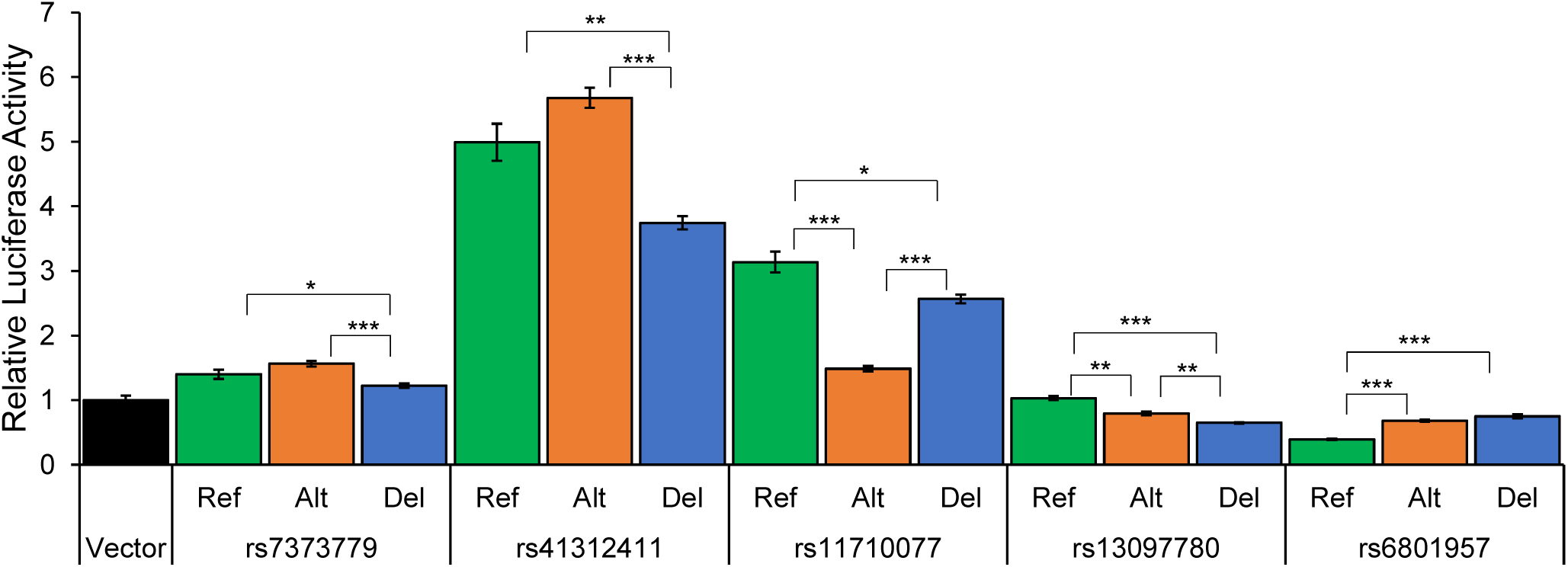
Firefly luciferase reporter enhancer assays in HL1 cells using variant-centered amplicons with reference (Ref), alternate (Alt) or 11-base deletion (Del) alleles for the five causal regulatory variants at the *SCN5A* QT interval GWAS locus. Bar plots show relative luciferase activities from empty vector and test constructs (firefly luciferase expression relative to *Renilla* luciferase expression and normalized to the expression from empty vector). Asterisks indicate *P* values from the Student’s *t* test (\**P*<0.05; \*\**P*<0.01; \*\*\**P*<0.001). Error bars are SEMs (*n*=4).

### Identification of putative TFs driving allele-specific enhancer activities

Given that the variant sites and their immediate flanking sequences are critical for driving enhancer activities at the five *SCN5A* enhancers (above), we next used DNA oligo pulldown assays to identify putative TFs that bind to the enhancer variants and their flanking sequences. First, variant-centered input sequences representing the reference and alternate alleles at each enhancer variant along with their flanking bases were scanned against a database of vertebrate TF position weight matrices (Dataset S2) to predict TF binding. Of the total 43 predictions, based on higher human cardiac (left ventricle) expression (29), 15 predicted TFs for four *SCN5A* enhancer variants (rs7373779, rs41312411, rs11710077 and rs6801957) were selected for binding assays. The two TFs predicted to bind rs13097780 had low cardiac expression and were not evaluated further. Given the *in vitro* nature of binding assay and the technical limitations of bacterial expression of full-length proteins, we expressed and purified the DNA-binding domains of the selected TFs as N-terminal GST-fusion proteins (Dataset S2). Streptavidin agarose beads-based pull down of biotinylated DNA oligos-TF complexes was evaluated by SDS-PAGE and Coomassie blue staining (Figure 2). At rs7373779-based oligo, GATA2 was predicted to bind the reference allele, but not the alternate allele, and that was observed in the pulldown assay as well. We did not observe binding of GATA4 or GATA6 at rs7373779-based oligo (data not shown). At rs41312411-based oligo, SP1, SP2, KLF4 and KLF10 were predicted to bind the reference allele, but not the alternate allele, and MAZ was predicted to bind both alleles. Pulldown assays showed SP1 binding to reference allele of rs41312411, but not the alternate allele as predicted. We did not observe binding of SP2, KLF4, KLF10 and MAZ at rs41312411-based oligo (data not shown). At rs11710077-based oligo, ERF, ELK1, ETV6 and SPI1 were predicted to the reference allele, but not the alternate allele. ERF, ELK1 and SPI1 binding to rs11710077 reference allele were supported by pulldown assays. However, in contrast to the *in silico* prediction, SPI1 was found to bind rs11710077 alternate allele as well. We did not observe binding of ETV6 at rs11710077-based oligo (data not shown). And at the rs6801957-based oligo, MEIS2 and MEIS3 were predicted to bind only at the alternate allele, and ZNF189 was predicted to bind only at the reference allele. In support of the predictions, MEIS2 and MEIS3 only bound the alternate allele at rs6801957, and ZNF189 was found to bind both alleles at rs6801957, but with a higher affinity for the reference allele. Together, based on *in silico* TF binding predictions and experimental confirmation by oligo-based pulldown assays, GATA2, SP1, ERF/ELK1 and MEIS2/MEIS3 binding likely drives the allele-specific enhancer activities at rs7373779, rs41312411, rs11710077 and rs6801957 enhancer variants, respectively.

**Figure 2:**
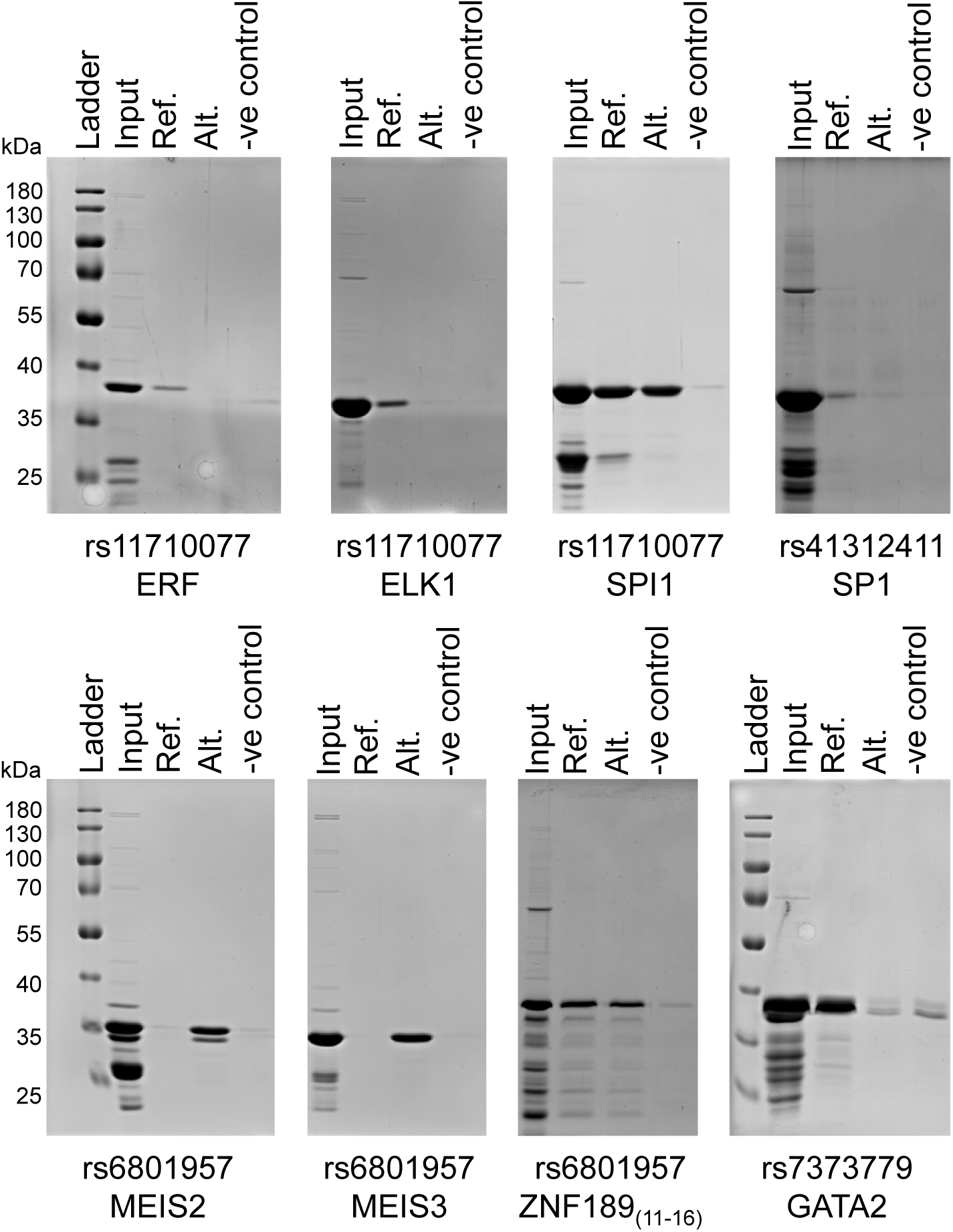
Oligo-based pulldown assays for TFs predicted to bind at rs7373779, rs41312411, rs11710077 and rs6801957 enhancer variants. Gel images show streptavidin agarose beads-based TF-GST fusion protein capture using biotinylated reference (Ref.) or alternate (Alt.) allele-based oligos, along with the TF-GST fusion protein alone (Input) and a pulldown without biotinylated oligo (-ve control; TF-GST fusion protein + streptavidin agarose beads) as controls. kDa: kilodaltons.

### Additive enhancer effects across the five SCN5A enhancer variants

Even though the five *SCN5A* enhancers centered on rs7373779, rs41312411, rs11710077, rs13097780 and rs6801957 were identified individually as enhancers with significant allelic differences using *in vitro* reporter assays, they were shown to collectively influence human cardiac *SCN5A* expression (8). In order to assess if the enhancer activities of the five enhancers are additive in nature, an expectation from their combined *cis* effects on gene expression, we evaluated enhancer activities of synthetic constructs made of the five elements in tandem (in genomic order) and in all possible allelic combinations using *in vitro* reporter assays. Thirty-two synthetic constructs representing all combination of the two alleles at each the five elements centered on rs7373779, rs41312411, rs11710077, rs13097780 and rs6801957 in tandem were cloned upstream of a minimal promoter-driven firefly luciferase reporter gene in pGL4.23 vector. Luciferase reporter assays were performed in mouse cardiomyocyte HL1 cells using all 32 synthetic constructs and empty vector control. Using the five variants, each with two allelic states (reference or alternate), as covariates in linear regression showed a significant effect (adjusted *R*^2^=0.65; *P*=3.07×10^-6^) on luciferase reporter activities from all the five variants (Supplemental Table S1). In this synthetic design, alternate alleles at each variant had a significant negative effect on overall enhancer activity except at rs6801957 where the direction of effect was opposite. Binning the 32 constructs based on the number of enhancer alleles in them (range: 0 to 5) indicated a clear additive effect on overall enhancer activity across the five enhancers (Figure 3).

**Figure 3:**
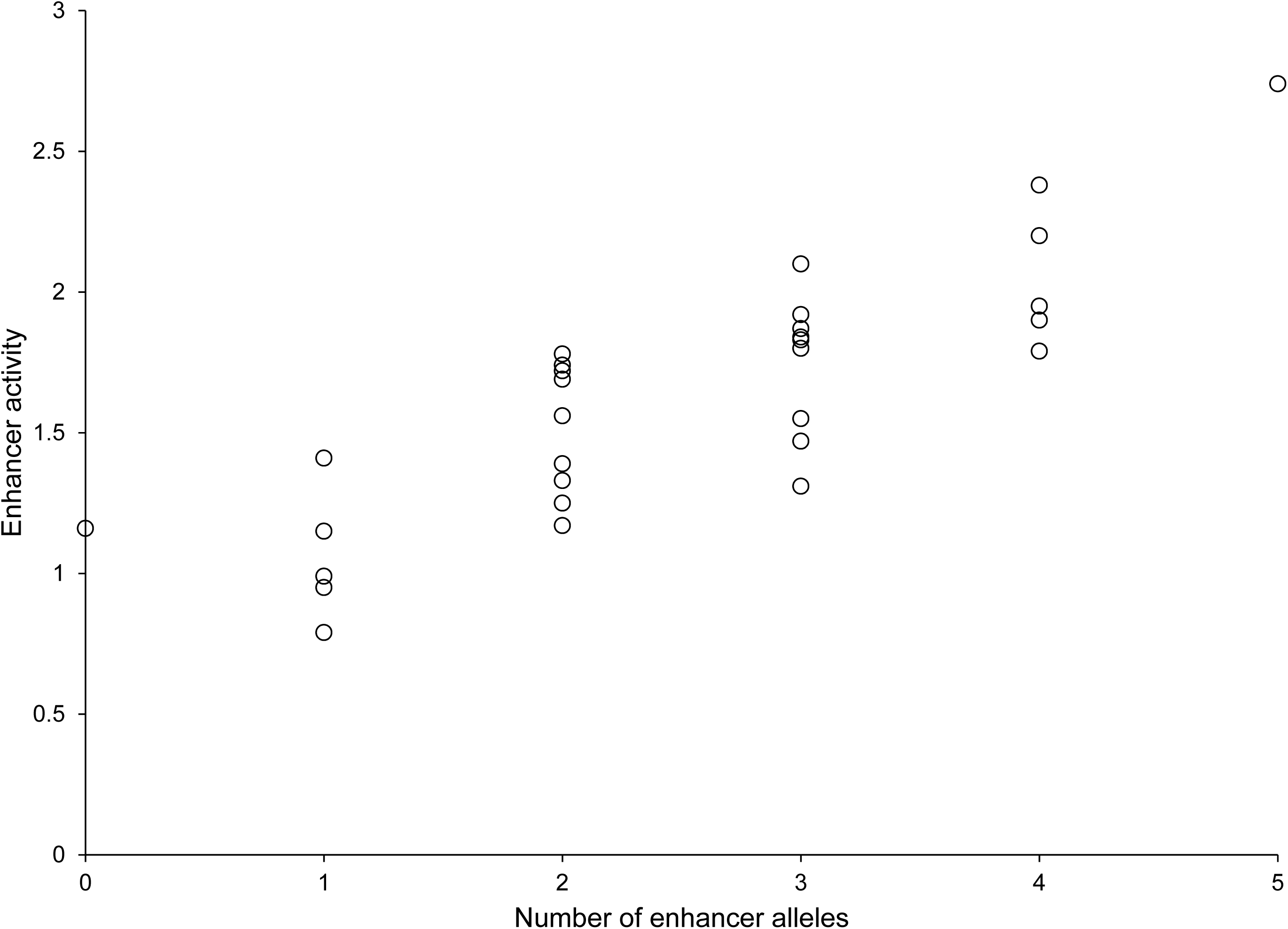
Combined additive effects from the five *SCN5A* biallelic enhancer elements on reporter expression. The scatter plot shows the enhancer activities measured by reporter assays in HL1 cells in comparison to the number of enhancer alleles per construct from the 32 synthetic constructs representing all possible allelic combinations across the five *SCN5A* biallelic enhancer elements.

### Four out of the five SCN5A enhancers function as enhancers in vivo

Having observed *in vitro* enhancer activities at the five *SCN5A* enhancers in mouse cardiomyocyte HL1 cells, we evaluated the potential of these elements to act as enhancers *in vivo* by using Tol2 transposase-mediated, random integration based transient developmental enhancer assays in zebrafish embryos. Variant-centered amplicons for rs7373779, rs41312411, rs11710077, rs13097780 and rs6801957 used in *in vitro* assays (above) were subcloned upstream of minimal promoter driven fluorescent reporter vectors (Dataset S1), injected into developing zebrafish embryos at 1-2 cell stage and observed for reporter expression 72hpf. Among the two alleles at each test element, reference and alternate, zebrafish enhancer assays were limited to the allele with higher activity in *in vitro* reporter assays. All except the rs13097780 centered element drove reporter expression in developing heart and somites (Figure 4, Supplemental Table S2), which overlaps with the endogenous expression of zebrafish homolog *scn5lab* in early development (30). As expected from transient assays, reporter expression for a given element was variable across injected embryos, and low levels of mosaic expression was also observed. Among the four elements with *in vivo* enhancer activities, the reporter expression was high for rs11710077 and rs6801957 centered elements (Figure 4).

**Figure 4:**
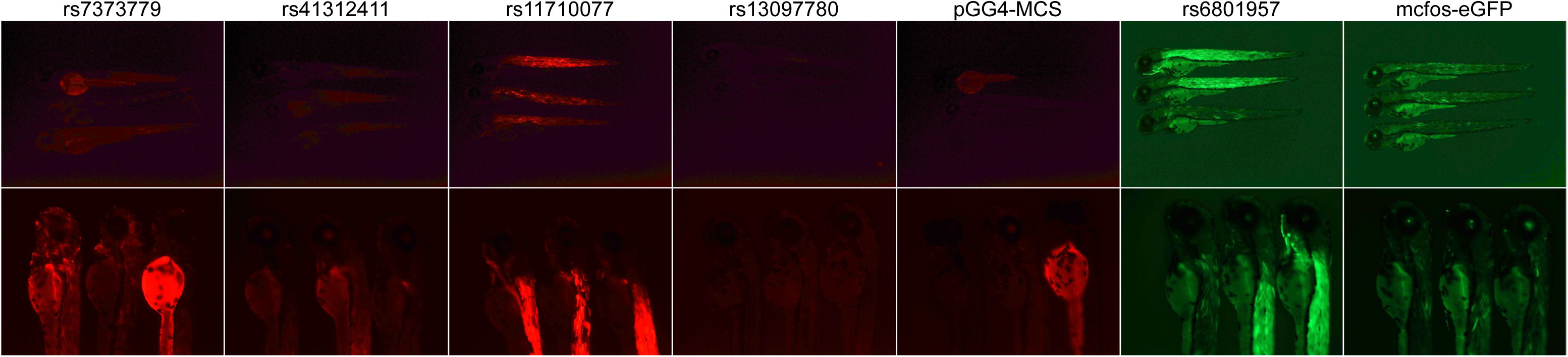
Transient enhancer assays in developing zebrafish embryos for the five *SCN5A* enhancer elements. Representative images of zebrafish embryos (entire body length (*top*) and zoomed in at the anterior part (*bottom*)) at 72hpf showing fluorescent reporter (RFP or GFP) expression. pGG4-MCS is the empty vector with RFP reporter for rs7373779, rs41312411, rs11710077 and rs13097780 centered elements, and mcfos-eGFP is the empty vector with GFP reporter for rs6801957 centered element.

## Discussion

Enhancers as *cis*-regulatory elements drive the transcriptional activity of target gene promoters, and multiple enhancers together lead to precise spatiotemporal regulation of gene expression (31). Although the phenomenon of enhancer cooperation has been investigated at the level of enhancers per se and individual genes, studies assessing the impacts from common regulatory variants, which disrupt individual enhancer activities on such cooperativity are lacking. Here, by functional characterization of QT interval associated five *SCN5A* enhancer variants and by evaluating all possible combinatorial constructs of these five biallelic elements, we demonstrate allele-specific additive effects on reporter gene expression *in vitro*.

We had previously reported (8) identification of five causal enhancer variants underlying the QT interval GWAS locus at *SCN5A*. To the best of our knowledge, it was the first study to report common regulatory variants at *SCN5A*, a gene known for its role in LQTS, impacting on QT interval variation in the general population. Collectively, the five enhancer variants were significantly correlated with human cardiac *SCN5A* expression and capable of explaining the QT interval associations (8). Nevertheless, which TFs likely drive allele-specific activities of enhancer variants and whether these elements act as *in vivo* enhancers remained unaddressed. Although expected from their combined *cis* effects on *SCN5A* cardiac expression, the nature of this combinatorial effect also remained unaddressed.

As *cis*-regulatory activity of enhancers is an outcome of concerted action of multiple TF binding sites (31), conclusions drawn from comparisons of allelic activities between reporter assays based on genomic elements and TF pulldown assays based on short oligos are not necessarily straightforward. Nevertheless, data from the two assays (Figures 1 and 2) can be used to build testable hypotheses. At rs7373779-centred enhancer element, enhancer activity (reporter expression) from the variant-centered 11bp deletion allele construct was reduced as compared to the reference and alternate allele constructs, which had similar activities, suggesting loss of a critical TF binding in the deletion allele. Based on oligo pulldown assays, GATA2 is the likely TF which binds at the reference allele construct and drives a higher enhancer activity as compared to the deletion allele construct, where the GATA2 binding site is lost. At the same time, enhancer activity from the alternate allele construct, where based on pulldown assays GATA2 is not expected to bind, is maintained at a level similar to that from the reference allele, thereby suggesting contributions from other TF bindings in the background. A similar principle can be applied at the rs41312411-centered enhancer element, where the reference allele construct, based on oligo pulldown assays, is bound by SP1, the deletion allele construct has relatively lower enhancer activity due loss of SP1 binding site, and enhancer activity in the alternate allele construct, likely not bound by SP1, is maintained by other TF bindings in the background. Based on enhancer assays for the rs11710077- and rs6801957-centered elements alone, one could conclude that the relatively higher enhancer activities from deletion allele constructs is due to removal of repressive TF binding sites present in the rs11710077 alternate allele and rs6801957 reference allele constructs. However, based on oligo pulldown assays, a more likely explanation for the observed enhancer activities at rs11710077-cenetred element is that rs11710077 reference allele construct is bound by ERF/ELK1 that leads to highest enhancer activity, alternate allele construct is not, and that the deletion allele creates a neo mutation (novel binding site) with intermediate effect. Similarly, based on oligo pulldown assays, a more like explanation for the observed enhancer activities at rs6801957-cenetred element is that rs6801957 alternate allele construct is bound by MEIS2/MEIS3 that leads to higher enhancer activity, reference allele construct is not, and that the deletion allele creates a novel binding site with an effect similar to that of the alternate allele.

Most enhancer discoveries are at individual enhancer level, whereas most genes are likely regulated by multiple enhancers. Therefore, understanding of the combined effects from multiple enhancers is essential to have a comprehensive understanding of gene expression regulation. A regression model to fit the observed variation in enhancer activities across the 32 possible allelic combinations of the five biallelic enhancer elements, clearly demonstrated additive effects across the five enhancers (Figure 3).

Although, at each of the five enhancers the allelic states, used a covariate in the linear regression, had a significant effect on overall enhancer activity, the allelic direction of the effect was inconsistent with that observed in single enhancer assays for two out of the five enhancer variants (Figure 1, Supplemental Table S1). These observations are not unexpected as enhancer activities of test elements (single- or multi-enhancer based) are dependent upon the context, which includes cooperative interactions between multiple TF binding sites across the entire length of the test element. Furthermore, it is important to note that though the additive interactions across these five biallelic enhancers are expected to explain a significant proportion of the variance in *SCN5A* cardiac expression at the population level, *SCN5A* expression variability is not entirely heritable and additional enhancers and their variants likely exist, beyond the ones underlying QT interval GWAS signal.

We have demonstrated that the majority of these *SCN5A* enhancers individually act as enhancers *in vivo* using transient enhancer assays in developing zebrafish (Figure 4), however, the additive effects described here are from *in vitro* studies using episomal constructs. Although, these assays do meet the classical definition of *cis*-regulatory enhancers (32), establishing their endogenous activities, individual as well as additive, in their native genomic context in human cardiomyocytes is warranted (33).

## Acknowledgements

This work was supported by funds from National Institutes of Health (NIH), USA Grant R01 HL158901 (to A.K.) and McGovern Medical School UTHealth (to A.K.). M.T.B. is supported by a Cancer Prevention and Research Institute of Texas Grant RP180804 for the Protein Array & Analysis Core.

## Author Contributions

A.K. conceived and designed the study. L.G., P.S., B.B., L.S., E.S., A.S., G.B., D.B. and A.K. conducted the experiments. L.G., P.S., B.B., M.T.B., D.B. and A.K. analyzed the experimental data. L.G. and A.K. wrote the manuscript. All authors were involved in manuscript revision.

