## Supplemental for "Functional characterization of QT interval associated *SCN5A* enhancer variants identify combined additive effects"

This PDF file includes:

Tables S1 to S2

Other supplementary materials for this manuscript include the following:

Dataset S1

Dataset S2

**Table S1: Effects of the alternate alleles at the five enhancer variants on overall enhancer activities observed in synthetic constructs representing all allelic combinations of the five enhancers in tandem**

| **Regression Statistics** | | | | | | |
|---|---|---|---|---|---|---|
| Multiple R | 0.84 | | | | | |
| R Square | 0.71 | | | | | |
| Adjusted R Square | 0.65 | | | | | |
| Standard Error | 0.25 | | | | | |
| Observations | 32 | | | | | |
| **ANOVA** | df | SS | MS | F | Significance F | |
| Regression | 5 | 3.97 | 0.79 | 12.49 | 3.07E-06 | |
| Residual | 26 | 1.65 | 0.06 | | | |
| Total | 31 | 5.63 | | | | |
| **Predictors** | Coefficients | Standard Error | t Stat | P-value | Lower 95% | Upper 95% |
| Intercept | 2.09 | 0.11 | 19.16 | 7.46E-17 | 1.87 | 2.32 |
| rs7373779 | -0.25 | 0.09 | -2.83 | 8.82E-03 | -0.44 | -0.07 |
| rs41312411 | -0.36 | 0.09 | -4.00 | 4.73E-04 | -0.54 | -0.17 |
| rs11710077 | -0.31 | 0.09 | -3.48 | 1.80E-03 | -0.49 | -0.13 |
| rs13097780 | -0.33 | 0.09 | -3.66 | 1.13E-03 | -0.51 | -0.14 |
| rs6801957 | 0.32 | 0.09 | 3.60 | 1.31E-03 | 0.14 | 0.50 |

### Table S2: Summary of microinjections for transient enhancer assays in developing zebrafish embryos

| Construct | # Injected | # Scored at 72hpf | # xFP+ at 72hpf | # Heart xFP+ 72hpf | % Survived | % xFP+ | % Heart xFP+ |
|---|---|---|---|---|---|---|---|
| rs7373779_pGG4-MCS | 275 | 167 | 140 | 46 | 60.73 | 83.83 | 32.86 |
| rs41312411_pGG4-MCS | 348 | 257 | 237 | 71 | 73.85 | 92.22 | 29.96 |
| rs11710077_pGG4-MCS | 331 | 218 | 200 | 102 | 65.86 | 91.74 | 51.00 |
| rs13097780_pGG4-MCS | 275 | 186 | 145 | 7 | 67.64 | 77.96 | 4.83 |
| rs6801957_pT2-mcfos-eGFP | 217 | 154 | 123 | 58 | 70.97 | 79.87 | 47.15 |
| pGG4-MCS | 381 | 275 | 251 | 8 | 72.18 | 91.27 | 3.19 |
| pT2-mcfos-eGFP | 285 | 174 | 106 | 36 | 61.05 | 60.92 | 33.96 |

xFP+: RFP or GFP positive.
